# Extending tactile space with hand-held tools: A re-analysis and review

**DOI:** 10.1101/2024.04.01.587537

**Authors:** Luke E. Miller, Alessandro Farnè

## Abstract

Tools can extend the sense of touch beyond the body, allowing the user to extract sensory information about distal objects in their environment. Though research on this topic has trickled in over the last few decades, little is known about the neurocomputational mechanisms of extended touch. In 2016, along with our late collaborator Vincent Hayward, we began a series of studies that attempted to fill this gap. We specifically focused on the ability to localize touch on the surface of a rod, as if it were part of the body. We have conducted eight behavioral experiments over the last several years, all of which have found that humans are incredibly accurate at tool-extended tactile localization. In the present article, we perform a model-driven reanalysis of these findings with an eye towards estimating the underlying parameters that map sensory input into spatial perception. This reanalysis revealed that users can almost perfectly localize touch on hand-held tools. This raises the question of how humans can be so good at localizing touch on an inert non-corporeal object. The remainder of the paper focuses on three aspects of this process that occupied much of our collaboration with Vincent: the mechanical information used by participants for localization; the speed by which the nervous system can transform this information into a spatial percept; and whether body-based computations are repurposed for tool-extended touch. In all, these studies underscore the special relationship between bodies and tools.

## Introduction

There is an intuitive sense—often evoked in literature (Butler, 1872)—that tools extend our body when they are used. This view is perhaps not surprising, given that tool use is a major part of our everyday lived experience as humans (Merleau-Ponty, 1962). In the early 20^th^ century, the British neurologists Henry Head and Gordon Holmes even noted that our bodily senses of location and movement extend with hand-held instruments (Head & Holmes, 1911). Though it took over eighty years for this claim to begin to be researched empirically (Maravita & Iriki, 2004; Martel, Cardinali, Roy, & Farne, 2016), numerous studies in the last three decades have found that tools influence numerous neurocognitive processes underlying perception (Canzoneri et al., 2013; Cardinali et al., 2011; Miller, Cawley-Bennett, Longo, & Saygin, 2017; Miller, Longo, & Saygin, 2014; Reed, Betz, Garza, & Roberts, 2010; Sposito, Bolognini, Vallar, & Maravita, 2012; Witt, Proffitt, & Epstein, 2005) and action (Bahmad et al., 2020; Berti & Frassinetti, 2000; Biggio, Bisio, Avanzino, Ruggeri, & Bove, 2020; Cardinali, Brozzoli, Finos, Roy, & Farnè, 2016; Cardinali et al., 2009; Farnè, Iriki, & Làdavas, 2005; Ganesh, Yoshioka, Osu, & Ikegami, 2014; Iriki, Tanaka, & Iwamura, 1996; Martel et al., 2019; Umiltà et al., 2008). However, most of this research has focused almost exclusively on the effects of *controlling* a tool (Johnson-Frey, 2004), ignoring the fact that tools also extend our ability to *sense* the environment (Vaught, Simpson, & Ryder, 1968).

Tools can extend the haptic sense of their users (Burton, 1993). A blind person using their cane to haptically perceive their environment is a classic example of this. However, sensing with a tool is not limited to edge cases such as this. Sensory information is ubiquitous during tool-use, and we almost certainly continuously process this information (perhaps unconsciously) to help guide our behavior. Though the number of studies is admittedly limited, there is evidence that tools can be used to sense the softness of an object (LaMotte, 2000), the texture of its surface (Hollins, Lorenz, & Harper, 2006; Klatzky, Lederman, Hamilton, Grindley, & Swendsen, 2003; Yoshioka, Bensmaia, Craig, & Hsiao, 2007), and even its distance from the body (Burton, 1992; Carello, Fitzpatrick, & Turvey, 1992; Giudice, Klatzky, Bennett, & Loomis, 2013; Saig, Gordon, Assa, Arieli, & Ahissar, 2012). There is even some early hints that sensorimotor transformations underlying tactile localization are shared between body and tools (Yamamoto & Kitazawa, 2001b; Yamamoto, Moizumi, & Kitazawa, 2005). Still, our knowledge of the mechanisms underlying tool-extended sensing remained limited. It is in the context of this research that we teamed up with Vincent Hayward to chart how tools extend the user’s tactile space beyond their body.

In 2016, we three began a series of studies that attempted to push extended touch to its limits (Miller et al., 2018). Our question was simple: How accurately can humans localize where the surface of a hand-held tool was touched? To address this question, we transposed common paradigms for reporting the perceived location of touch on the body to measure the perceived location of touch on a tool (a wooden rod). Specifically, our participants pointed on a drawing of a rod where they felt the rod was touched. Touch was applied two ways: In a *passive* condition, participants localized touch applied passively to the rod surface; in an *active* condition, participants wielded the rod onto an object and judged where on the rod’s surface the strike occurred. The purpose of these manipulations was to assess whether movement (and presumably efference copies) were necessary for tool-extended perception, or whether sensory information was sufficient (the passive condition).

While we expected that localization would be possible, we did not expect how well. To our surprise, we found that humans could localize touch on a hand-held rod extremely accurately. This was the case when touch was passively applied to its surface and when the rod was actively used to contact the object, suggesting that sensory information alone is sufficient (to some extent) for localization. However, only when touch was active did we see near-perfect performance, highlighting tool-extended sensing as a closed-loop sensorimotor behavior (Ahissar & Assa, 2016). We have since performed numerous studies (Fabio, Salemme, Farne, & Miller, 2024; Fabio, Salemme, Koun, Farne, & Miller, 2022; Miller, Fabio, et al., 2023; Miller et al., 2019; Miller, Jarto, & Medendorp, 2023), both published and unpublished. These include behavioral studies, computational studies, neural network modelling, and neural recordings. In all, we find that humans are extremely accurate at localizing where a tool has been touched.

In memory of and dedication to Vincent—our dear friend and long-time collaborator— the time is ripe for us to synthesize the behavioral findings of all experiments we have conducted since our collaboration began. In total, we will synthesize the findings of eight experiments, both published and unpublished. In doing so, we aim to draw broader conclusions about the ability that would not be possible with a single experiment. Using a model-based approach (Miller et al., 2018), we address three aspects of the phenomenon: 1) Identifying the function that relates sensory input to the perception of touch location; 2) Characterizing the parameters of this function, and how they relate to perceptual accuracy; and 3) Determining the best task for accurately measuring tool-extended sensing.

We will then discuss the sensorimotor mechanisms underlying tool-extended sensing, focusing on the mechanical information used for sensing, the speed by which the nervous system processes this information, and the possibility that the brain repurposes body-based computations to sense with a tool. We conclude that during sensing, the tool should be viewed as an extended sensory ‘organ’ that is integrated within the somatosensory system.

## Methods

### Database

The re-analysis includes eight different datasets, six of which have been previously published (Miller, Fabio, et al., 2023; Miller, Jarto, et al., 2023; Miller et al., 2018). All experiments followed a similar experimental procedure, though there were relatively minor variations on specific aspects of how the task was implemented: i.e., method for reporting judgments, geometry of the rod etc.; These can be seen in Table 1.

**Table 1.** Details of each dataset used in the re-analysis.

| E | N | Judgment type | # Locs | Rod Material | Length | Reference |
| --- | --- | --- | --- | --- | --- | --- |
| 1 | 10 | Drawing | 7 | Wood | 83 (cm) | Miller et al 2018; E1 |
| 2 | 10 | Drawing | 7 | Wood | 83 | Miller et al 2018; E2 |
| 3 | 20 | Drawing | 6 | Wood | 60 | Miller et al 2018; E4 |
| 4 | 12 | Drawing | 6 | Wood | 60 | Unpublished |
| 5 | 10 | Drawing | 7 | Wood | 60 | Miller et al 2018; E5 |
| 6 | 38 | Drawing & Space | 6 | Wood & PVC | 60 | Miller et al., 2023 |
| 7 | 20 | Verbal report | 6-11 | Carbon fibre | 530-600 | Miller et al., 2023 |
| 8 | 37 | Physical model | 7 | Wood | 80 | Unpublished |

### Participant Information

One-hundred and fifty-seven participants in total took part in the experiments. All participants had normal or corrected-to-normal vision, and were free from neurological and sensorimotor conditions. Informed consent was taken before the start of the experiment. We refer the reader to the original studies for greater demographic details.

### Experimental methods

In every experiment, the task of the participant was to localize touch on a hand-held rod. The details for each dataset can be seen in Table 1; for an in-depth treatment of the methods, we refer the reader to each paper. We will first describe the general structure that is shared across participants. We will then briefly discuss the different reporting methods used across experiments.

The general experimental structure was as follows: Participants sat comfortably at a table, holding the rod out-of-view behind an occluding board. At the beginning of each trial, the participant was instructed to hold the rod still (pre-contact phase). Upon receiving a cue, they actively wielded the rod downward to contact an object (contact phase), which was placed on the table at a variable distance from the participant’s hand. After contact, the participant reported where on the tool they judged the touch to have occurred (see below). The object was typically placed in 6–7 unique locations, with 10–20 trials per location. The localization task typically took ∼15–30 minutes to complete, depending on the specific experiment.

The method that they reported their judgment varied per experiment. In the *Drawing task* (E1–6), participants used a computer cursor to report on a drawing of the rod where they localized the touch. In the *Space task* (E6), participants used a computer cursor to point to the location in an empty screen that corresponded to where they localized the touch. In the *Verbal report task* (E7), participants verbalized where they felt the touch, from 0–100 (base–tip). In the *Physical model task* (E8), participants pointed on a physical model of the tool to the corresponding touch location. With each of these tasks, we aimed to determine how accurate each participant could localize touch on the tool.

### Data analyses

#### Model Fitting

As in Miller et al., (2018), we performed a model-based analysis to characterize the perceptual function underlying each participant’s localization behavior. We considered two models for how a tool-user might localize touch on the tool. They are discussed below:

##### Extension Model

Here, the perception of touch is extended along the entire surface of the rod. There is therefore a linear relationship between the judged and the actual location of touch. We can model each participant’s behavior with a linear regression:

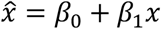

where the two free parameters are the offset (*β*_0_) and the slope (*β*_1_). The offset corresponds to a general bias towards or away from the hand. The slope corresponds to the gain in the judged location 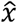 as a function of the actual touch location *x*. Both the offset and slope were free parameters in the model fitting. Note that this model was called the *Embodiment Model* in our original article.

##### Funneling Model

Here, the perception of touch is funneled towards one of the extreme ends of the rod (handle or tip). In its most extreme form, the perceived location of touch in the first half of the rod is funneled completely to the handle and touch in the second half of the rod completely to the tip. There is therefore a sigmoidal relationship between the judged and the actual location of touch. We can model each participant’s behavior with the following equation:

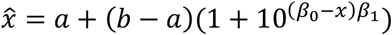

where the four parameters are the minimum (*a*) and maximum (*b*) of the curve, the midpoint (*β*_0_) and the slope (*β*_1_). All four parameters were set to free in the model fitting. Note that this model was called the *Projection Model* in our original article.

#### Model Comparison

We used the Bayesian Information Criterion (BIC) to compare the ability of the Extended and Funneling Model to fit each participant’s data.

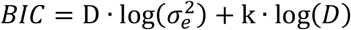

Where *D* is the number of datapoints, *k* is the number of free parameters, and 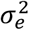 is the mean squared error of the fit. The lower the BIC score, the better the model fit. We can therefore use the difference between each model’s BIC score (Δ*BIC*) as a means for determining whether one model significantly fit the data better than the other. A Δ*BIC* > 3 corresponds to a significantly better fit for the Extended Model compared to the Funneling Model. A Δ*BIC* < −3 corresponds to a significantly better fit for the Funneling Model compared to the Extended Model.

## Results

### Localization behavior is consistent with the extension of touch

We first sought to compare each model’s ability to fit each participant’s dataset. As in our original article, the Extension Model was by far the best fitting model (Figure 1A; mean±sem; ΔBIC: 12.77±0.76). This was particularly apparent in the individual participant fits. Out of 157 total participants, the Extension Model was the significantly better model (ΔBIC>3) in 143 participants; 138 of those showed strong evidence (ΔBIC>6) for the Extension Model (Figure 1B). Only six participants showed significantly better evidence for the Funneling Model.

**Figure 1.**
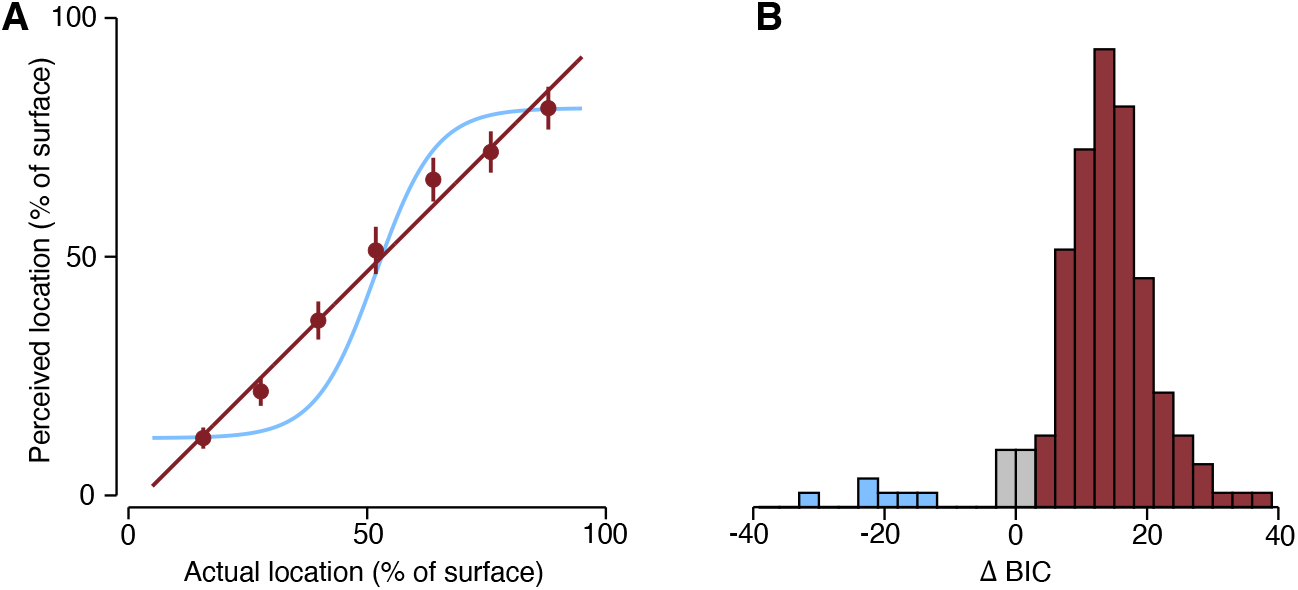
Results of model-based analysis. (A) The fit to the data from Miller et al., (2018) with the Extended model (red) and Funneling Model (blue). (B) The delta-BIC for all participants, whose dataset favored extension (red), funneling (blue), or neither (gray).

In its most extreme form, the Extension Model predicts a unity between the actual and perceived location of touch (i.e., intercept of zero; slope of one). This model also fit the data better than the Funneling Model (Figure 1A; mean±sem; ΔBIC: 12.59±0.77). Out of 157 total participants, the Strong Extension Model was the significantly better model in 141 participants, with 132 of those showing strong evidence (ΔBIC>6) in favor. Only ten participants showed significantly better evidence for the Funneling Model.

### General parameter estimation

Having established that the behavior of our participants was best explained by the Extension Model, we next aimed to estimate the underlying parameters in order to quantify the accuracy of localization. The model fits for each dataset can be seen in Figure 2.

**Figure 2.**
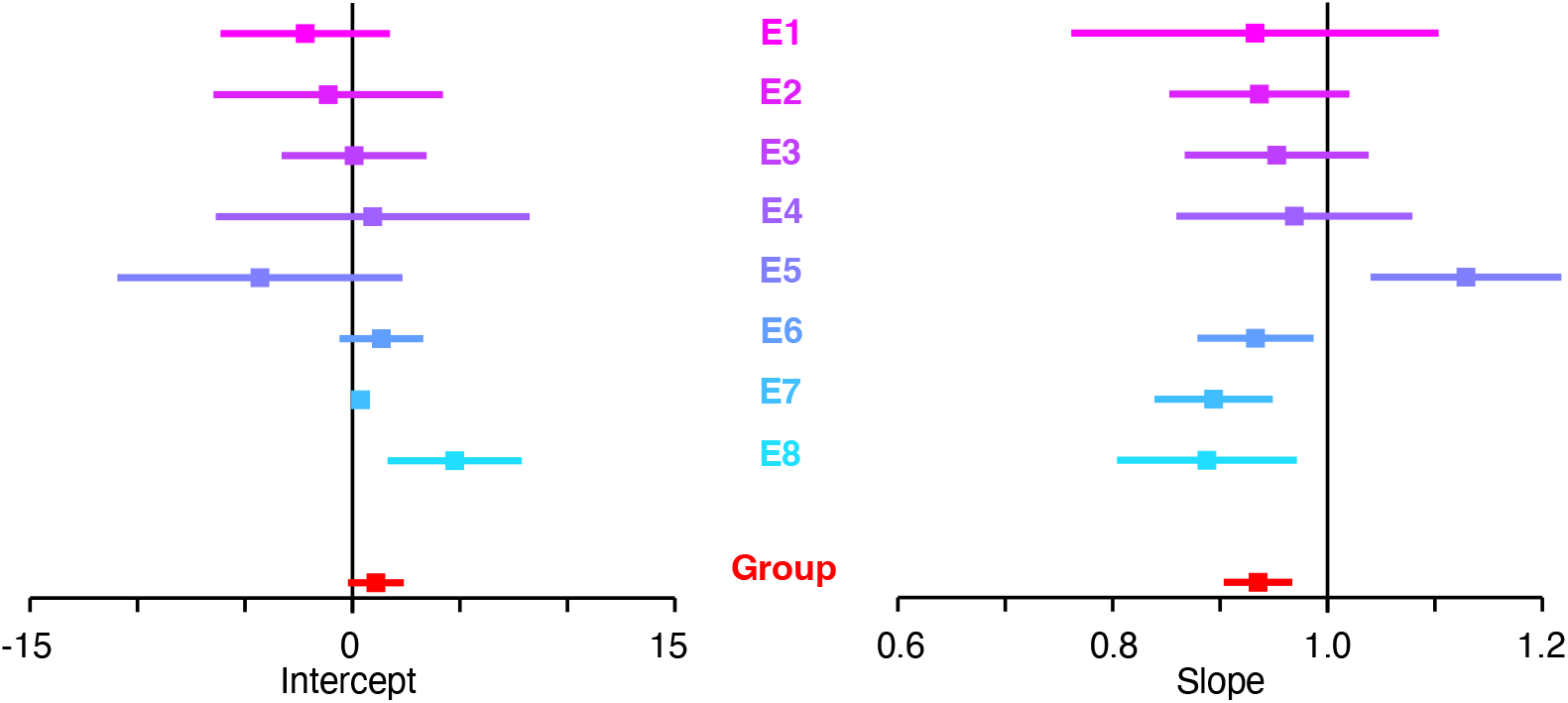
Parameter estimates for all experiments. Forrest plot of the mean estimates for each experiment (top–bottom; E1–E8) for the intercept (left) and slope (right) from the linear model. The red corresponds to the group aggregated estimate. Error bars correspond to the 95% confidence interval of the estimate.

The intercept quantifies a general localization bias (e.g., towards the hand). Across all datasets (Figure 2A), the mean intercept ranged from -4.30 to 4.75 (in units of % of surface). The confidence intervals of the estimates in each experiment nearly always including zero. Indeed, the group-level estimate was 1.09 (95% CI [-0.21; 2.39]), demonstrating that localization is characterized by a small-to-nonexistent general spatial bias away from the hand in judgments.

The slope is the most informative estimate for our purposes, as it quantifies the correspondence between where the tool was touched and where the participant felt it was touched (i.e., localization accuracy). Localization accuracy was consistently high across all datasets (Figure 3B), with the observed mean slopes ranging from 0.89 to 1.13 (in units of % of surface). The confidence intervals of the estimates in each experiment nearly always including one (i.e., unity). The group-level estimate (mean: 0.94; 95% CI [0.90; 0.97]) demonstrated that while localization was significantly different than unity, it was extremely high. In general, we can conclude that localization was highly accurate across all eight datasets (Table 2).

**Table 2.** Parameter estimation for each dataset.

| Exp | Judgment | Intercept |  | Slope |  |
| --- | --- | --- | --- | --- | --- |
|  |  | Mean | 95% CI | Mean | 95% CI |
| 1 | Drawing | -2.20 | [-6.14; 1.74] | 0.93 | [0.76; 1.10] |
| 2 | Drawing | -1.13 | [-6.47; 4.21] | 0.94 | [0.85; 1.02] |
| 3 | Drawing | 0.08 | [-3.30; 3.45] | 0.95 | [0.87; 1.04] |
| 4 | Drawing | 0.94 | [-6.36; 8.25] | 0.97 | [0.86; 1.08] |
| 5 | Drawing | -4.30 | [-10.94; 2.33] | 1.13 | [1.04; 1.21] |
| 6 | Drawing | 1.34 | [-0.60; 3.29] | 0.93 | [0.88; 0.99] |
|  | External space | 5.43 | [2.71; 8.15] | 0.89 | [0.82; 0.96] |
| 7 | Verbal response | 0.38 | [0.16; 0.60] | 0.89 | [0.84; 0.94] |
| 8 | Physical Model | 4.75 | [1.63; 7.88] | 0.88 | [0.80; 0.96] |
| Group |  | 1.09 | [-0.21; 2.39] | 0.94 | [0.90; 0.97] |

**Figure 3.**
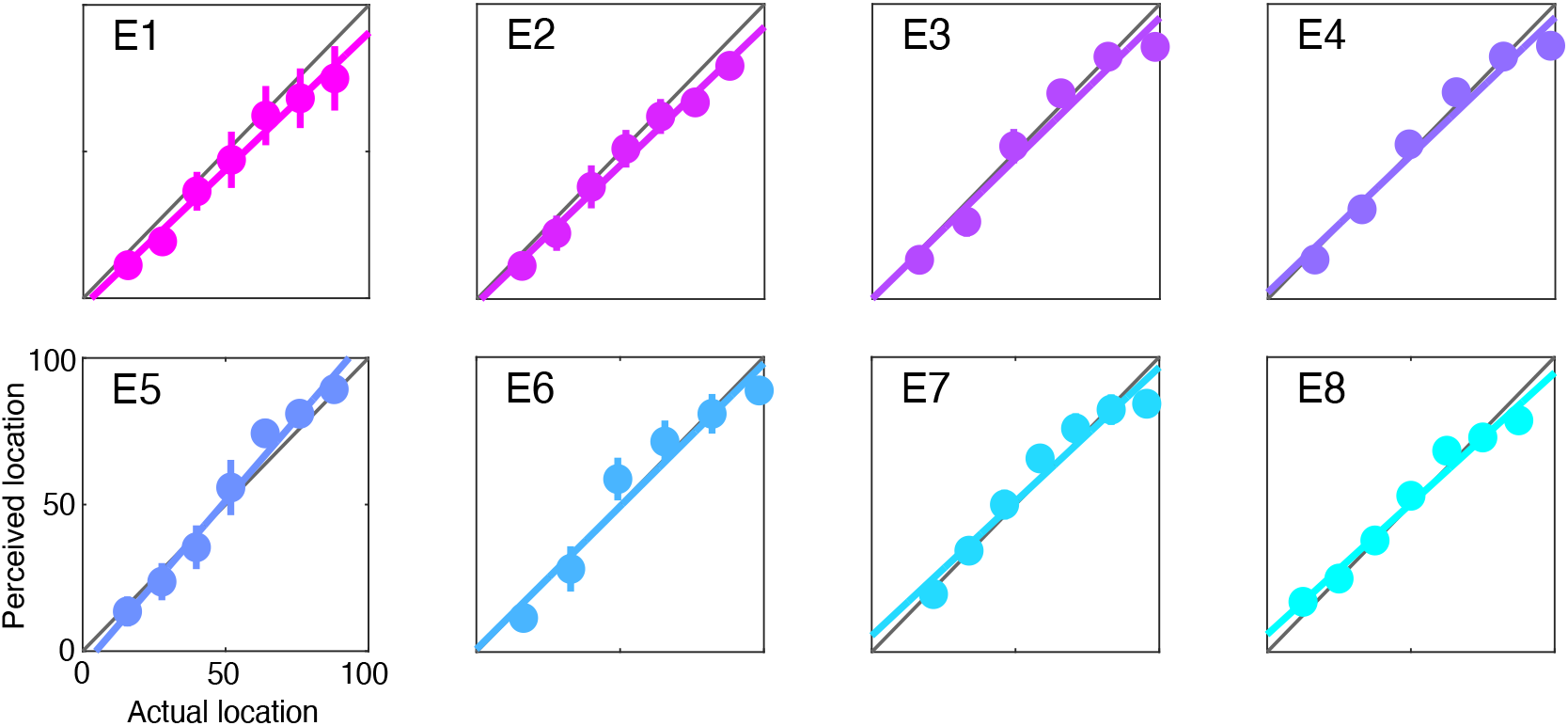
Model fits for each experiment. The fit of the linear model to each experiment (E1–E8), with perceived location modelled as a function of actual location (in units % of surface). Error bars over the mean judgments correspond to the 95% confidence interval.

### The effect of reporting type on accuracy estimation

We next took a more fine-grained look at how the estimation of localization accuracy varied across task. We assume that participant’s internal localization accuracy does not depend upon the required method of reporting. Instead, our measurement of the judgments reported with any method merely reflect a noisy readout of the participant’s internal estimate of touch location. The accuracy estimate we measure in each experiment should therefore be thought of as a lower-bound on how accurate that internal estimate of location was. It is therefore beneficial to use a method of reporting that will give you the most accurate estimate of localization.

Across all eight datasets, we used four different reporting methods to measure localization judgments: participants reported their judgments on a drawing of a rod (n=100; E1–6), with a cursor in external space (n=38; E6), on a physical model of a rod (n=37; E8), and verbally (n=20; E7). The drawing task demonstrated that best correspondence between judgments and actual touch location, with a mean slope of 0.96 and an upper bound on the confidence interval a at unity (95% CI [0.92; 1.00]). Though the estimate of accuracy in the other tasks was substantially worse (mean slopes: 0.89), the upper bound on the confidence intervals still approached unity (see, Table 2). In general, these findings demonstrate that the drawing task provided the most accurate read-out of participant’s localization abilities.

## Discussion

Individually, all eight studies demonstrated that humans can accurately localize touch on hand-held rods—even when the rod is over five-meters long. In the present study, we aimed to go beyond individual datasets and re-analyze them all collectively. In doing so, our main goals were to characterize the type of sensing done by tool-users as well as the parameters underlying their perception. We can conclude three things about tool-extended sensing from our re-analysis: First, the localization patterns of nearly all 157 participants matched what would be expected had they linearly extended tactile perception beyond the body and onto the rod. Second, this ability was extremely accurate, with a global bias near zero and a linear gain approaching one. Third, and relatedly, the measured localization accuracy depended on the type of reporting task used in the experiment. Participants displayed the most accurate judgments when reporting touch location on a drawing. Since behavioral performance reflects a lower bound on perceptual accuracy, we can conclude from the results of this task that localization accuracy is nearly perfect; that is, the confidence interval for the slope included one (i.e., unity).

The above results are highly suggestive that when sensing with a tool, it becomes integrated with the sensorimotor system to function like an extended ‘sensor’. However, they are not the only pieces of evidence that led us to conclude that tools extend the somatosensory system. Together with Vincent Hayward, we focused on three questions that round out the phenomenon of tool-extended sensing: 1) What mechanical information is used to extract objection location information with a rod? 2) How quickly is this information does the nervous system extract this information? 3) Does localization involve repurposing neurocomputational mechanisms that typically localize touch on the body? We focus on these three questions in the Discussion section.

### Mechanical determinants of touch location

Tactile localization starts in the periphery. When an object touches the body, its location is encoded by a specific subset of mechanoreceptors directly under its skin location (Johnson, 2001). This obviously cannot be the case for touch on tools since they are not innervated. This is a major difference between body parts and non-body implements, and it raises the question of how exactly the localization process starts. Together with Vincent, we proposed the touch location is encoded in the mechanical response of the rod when contacting the object (Miller et al., 2018)—Specifically, the vibrations resulting from tool-object contact.

When a rod is struck, it resonates at specific frequencies called *modes*. For wooden rods, a subset of these modes are within 20–1000 Hz, well within the range of mechanoreceptors in the forelimb (Johansson & Flanagan, 2009). Importantly, the amplitudes of these modes depend upon *where* the strike occurred on the rod’s, a relationship that is formalized by the Euler-Bernoulli theory of beams. Therefore, striking a rod produces a location-specific pattern of vibrations that we termed a *vibratory motif*. What makes vibratory motifs such an intriguing potential cue used for sensing is that this location-amplitude relationship is *invariant* across most types of rods used by humans. Whereas the material and geometric properties of the rod determine the frequencies it will resonate at, the location-specific pattern is theoretically always the same. The location-amplitude dynamics are therefore learnable and could be implemented by internal models (Taunyazov et al., 2021).

We tested this proposal in an experiment with a hybrid tool that was made of two materials: the rod’s handle and half of its body were wood whereas the other half of the body was foam. Because participants never saw or used the tool prior to the experiment, once they grasped the tool they assumed it was entirely wood. When contact was made on the wooden portion, participants showed the same level of accuracy we observed in the previous experiments. However, when contact was made on the foam portion, localization proved impossible. Having tuned their internal model to anticipate the dynamics from a wooden rod they were unable to interpret the sensory signals when contact occurred on the foam half.

One interesting application of these findings is the field of prosthetics (Bensmaia & Miller, 2014), where restoring touch has been incredibly important (Raspopovic, Valle, & Petrini, 2021). Typically, this involves sensorizing the prosthesis and using the sensor readout to drive tactors (Kuiken, Marasco, Lock, Harden, & Dewald, 2007; Marasco, Kim, Colgate, Peshkin, & Kuiken, 2011), nerve stimulation (George et al., 2019), or neural implants (Flesher et al., 2021). This allows the prosthesis to feel more like the actual body part that was lost, improving object manipulation and embodiment. Given the exquisite sensitivity to the information carried in the intrinsic dynamics of a tool, we hypothesized that the intrinsic dynamics of a prosthesis might be picked up by wearers, allowing them to feel tactile information (Miller et al., 2019). Indeed, a recent study found that for upper limb prosthetic devices, the mechanical response of the prosthesis when touching an object does contain spatial information (Ivani et al., 2024). Most importantly, the wearer can indeed perceptually distinguish types of information from the intrinsic dynamics alone. As we proposed, leveraging the intrinsic mechanical response of a prosthesis could open new avenues for restoring touch.

### The speed of spatial coding during tool-extended sensing

During sensing, it is important that the information encoded by a sensor is rapidly extracted by the nervous system. For this reason, understanding the temporal aspects of tool-extended sensing was a major part in our investigations. Inspired by our theory of vibratory motifs, we first looked at how quickly location-specific information emerges in vibrations during sensing. We recorded vibrations in three participants while they performed our tool-sensing task (Miller et al., 2018). We then used machine learning to decode touch location (and perceived location) from the vibrations. By varying the amount of vibration information we supplied the decoder, we found that only ∼10 milliseconds of vibration are sufficient to completely distinguish the location of touch. Thus, vibrations are a perceptual cue that could be used for sensing in real time.

We next estimated how quickly this information could be re-encoded by mechanoreceptors in the hand. To do so, we used a biologically plausible skin-neuron model (Saal, Delhaye, Rayhaun, & Bensmaia, 2017) that can generate plausible spiking patterns given a time-varying input such as vibrations. We focused our modelling effort on Pacinian corpuscles, as these have broad frequency tuning in the range of the aforementioned resonant modes (Bell, Bolanowski, & Holmes, 1994), precise temporal spiking (Mackevicius, Best, Saal, & Bensmaia, 2012), and have been implicated in tool-use (Johnson, 2001). Combining machine learning with signal processing, we found that location information was encoded in the precise spike timing of the PC response. Furthermore, only ∼20 ms of spikes were necessary to decode hit-location with near 100% accuracy. Thus, where a tool has been touched emerges rapidly in pre-afferent and afferent codes.

We next recorded electroencephalography (EEG) during tool-sensing to address how quickly location-specific information emerges in neural dynamics (Miller et al., 2019). If the emergence of spatial coding is temporally efficient, it should emerge almost immediately after touch on the tool. The above afferent simulations provide a temporal lower-bound (TLB) on the emergence of fine-grained spatial coding. Given the 20 ms temporal delay between touch the response of primary somatosensory cortex (S1), we set the TLB on spatial coding to be ∼40 ms.

We used a repetition suppression paradigm to pinpoint location-specific somatosensory brain responses, and analyzed our data according to whether touch was in the same or different location as the previous touch. We observed significant suppression over contralateral sensorimotor channels within ∼50 ms, only 10 ms later than the TLB. Source reconstruction pinpointed these brain responses to the hand region of S1, suggesting that the vibration-to-space transformation occurs incredibly rapidly (perhaps even before S1). Neural network modelling suggested that this may only require a single processing step (Miller, Fabio, et al., 2023). Though we still do not know the actual computations that perform the vibration-to-space transformation, our series of study suggest that they are implemented rapidly enough for dynamic real-time sensing.

### Repurposing body-based computations for tool-extended sensing

A major aspect of our research programs has focused on the relationship between technology and the body. On the one hand, the peripheral encoding of touch location between body and tool is quite different (see above). On the other hand, there appear to be similarities in terms of localization performance and perhaps even neural processing (Gallivan, McLean, Valyear, & Culham, 2013; Iriki et al., 1996; Pazen et al., 2020; Umilta et al., 2008). It has been our opinion that these similarities reflect something deep about the nature of sensorimotor representation in the brain. A major component of our research with Vincent was therefore devoted to using computational approaches to characterize this similarity, focusing on the transformations underlying tactile spatial coding. There is now substantial evidence that body-based somatosensory computations are indeed repurposed (to some extent) to sense with tools (Fabio et al., 2024; Kilteni & Ehrsson, 2017; Miller, Fabio, et al., 2023; Pazen et al., 2020; Yamamoto & Kitazawa, 2001b; Yamamoto et al., 2005). We review this evidence below.

There is behavioral evidence that somatosensory computations for sensing with the body are repurposed to sense with a tool. Deriving a limb-centered spatial code for touch on a body part amounts to computing its distance from the body-part boundaries (e.g., the joints), a computation we call trilateration (Miller et al., 2022). We recently found that the same computation is used to localize touch within the tool (Miller, Fabio, et al., 2023). That is, touch location within the tool is determined by computing its distance from the handle and tip of the rod. Because there is a direct mapping between vibrations and tool space, trilateration can occur over the vibratory inputs to derive a spatial representation of where the tool was touched. This study—and others—raise the question of whether neural processes for localizing touch on the body are also repurposed to localize touch on a tool.

Initial evidence that body-based neural processes are repurposed for tool-sensing comes from the EEG study we discussed above (Miller et al., 2019). For most participants, we measured neural dynamics for both touch on a rod *and* touch on their arm. The time-course, scalp topography, and neural sources (S1 and PPC) of space-related brain responses was nearly identical in both cases. Crucially, using several different machine learning methods, we demonstrated that these similarities were not just superficial. For example, machine learning classifiers trained to decode the location of touch on the arm could be used to decode touch on the tool (and vice versa). This was true for the earliest brain responses (50 ms) that were localized to neural dynamics in S1. This cross-surface decoding demonstrates that these early brain responses are not just related to vibrations, as some have mistakenly claimed (Schone, Mor, Baker, & Makin, 2021). Instead, they reflect shared neural processes for the low-level spatial processing of touch on the body and on tools.

Acting towards a touch on the body requires remapping a low-level skin-based representation into a representation of touch location in external space (Heed, Buchholz, Engel, & Roder, 2015). Tactile remapping is often measured using a tactile temporal order judgment paradigm (Azanon, Mihaljevic, & Longo, 2016; Badde, Heed, & Roder, 2016; Yamamoto & Kitazawa, 2001a), where task performance decreases when the hands are crossed (crossed-hands deficit). This is likely due to spatial conflicts between multiple concurrent reference frames, e.g., when the *left* hand is placed in *right* external space (i.e., crossed). Interestingly, a few studies have also found evidence for a crossed-sticks deficit (Yamamoto & Kitazawa, 2001b; Yamamoto et al., 2005). When participants crossed two sticks across their midline, their ability to report which stick was touched first diminished, even though their hands remained uncrossed. This is suggestive of similar sensorimotor transformations remap touch on hands and tools in external space (Miyazaki, Hirashima, & Nozaki, 2010).

We recently investigated whether the neural correlates for tactile remapping on the hands are repurposed to remapping touch on a rod. Different tactile representations engage distinct neural oscillatory channels. Beta oscillations (15–25 Hz) have been implicated in localizing touch in skin-based coordinates (Buchholz, Jensen, & Medendorp, 2011), whereas alpha oscillations (8–13 Hz) map the location of touch in external space (Buchholz et al., 2011; Schubert et al., 2019). We recorded EEG while participants localized touch on uncrossed/crossed hands or uncrossed/crossed rods. In the latter, it is critical to note that hand posture remained unchanged in each condition—only the posture of the tools changed. Regardless, we found that the neural processes underlying tactile remapping on hands and tools were remarkably similar. Touch on hands and tools led to virtually identical space-related alpha band modulations throughout parietal cortex. These findings provide further evidence that localizing touch on tools involves repurposing body-based mechanisms for mapping touch on the body.

## Conclusion

We would like to conclude our paper with some reflection on our time with Vincent. He was instrumental to all discoveries, both empirical and theoretical, and helped shape the direction of the research we continue to this day. Given our history of studying embodiment, we can into the project viewing the phenomenon of tool-sensing from a purely psychological perspective. With Vincent’s help, we began to see the whole picture. To understand tool-extended sensing, we needed a wider lens—one that encompassed all levels of information transformation, from mechanics to neural computation. Vincent’s contribution to our thinking—to the widened lens by which we now view the sensorimotor world—was immense; the breadth of the two papers co-authored with him are a testament to this. We will be forever grateful and lucky to have had him in our intellectual and personal lives.

